# Pre-vaccination Frequency of Circulatory Tfh is associated with Robust Immune Response to TV003 Dengue Vaccine

**DOI:** 10.1101/2021.08.19.456926

**Authors:** Abdullah M Izmirly, Adam-Nicolas Pelletier, Jennifer Connors, Bhavani Taramangalam, Sawsan O. Alturki, Emma A. Gordon, Sana O. Alturki, Joshua C. Mell, Gokul Swaminathan, Vivin Karthik, Michele A. Kutzler, Esper G. Kallas, Rafick-Pierre Sekaly, Elias K Haddad

**Author notes:** Department of Medicine, Drexel University College of Medicine, Philadelphia, PA, United States of America.

## Abstract

It has been estimated that more than 390 million people are infected with Dengue virus every year; around 96 millions of these infections result in clinical pathologies. To date, there is only one licensed viral vector-based Dengue virus vaccine CYD-TDV approved for use in dengue endemic areas. While initially approved for administration independent of serostatus, the current guidance only recommends the use of this vaccine for seropositive individuals. Therefore, there is a critical need for investigating the influence of Dengue virus serostatus and immunological mechanisms that influence vaccine outcome. Here, we provide comprehensive evaluation of sero-status and host immune factors that correlate with robust immune responses to a Dengue virus vector based tetravalent vaccine (TV003) in a Phase II clinical cohort of human participants. We observed that sero-positive individuals demonstrate a much stronger immune response to the TV003 vaccine. Our multi-layered immune profiling revealed that sero-positive subjects have increased baseline/pre-vaccination frequencies of circulating T follicular helper (cTfh) cells and the Tfh related chemokine CXCL13/BLC. Importantly, this baseline/pre-vaccination cTfh profile correlated with the vaccinees’ ability to launch neutralizing antibody response against all four sero-types of Dengue virus, an important endpoint for Dengue vaccine clinical trials. Overall, we provide novel insights into the favorable cTfh related immune status that persists in Dengue virus sero-positive individuals that correlate with their ability to mount robust vaccine specific immune responses. Such detailed interrogation of cTfh cell biology in the context of clinical vaccinology will help uncover mechanisms and targets for favorable immuno-modulatory agents.

**Author summary:** Dengue virus (DENV) is a worldwide threat that causes significant health and economic burden. Currently, there are several challenges in the development of a DENV vaccine including the existence of four different serotypes all; capable of causing disease and antibody dependent enhancement (ADE). For complete protection, a vaccine must be able to generate neutralizing antibodies against all 4 serotypes to avoid ADE. Currently, there is one licensed DENV vaccine, CYD-TDV (DENGVAXIA^TM^). However, this vaccine is only efficacious in protecting against severe disease in DENV seropositive individuals therefore serostatus effect must be further studied for optimal vaccine design. A subset of CD4+ T cells called T-follicular helper (Tfh) cells have been well known to play a major role in aiding high affinity antibody production. Therefore, we chose to look at subsets of Tfh and the cytokines they produce in human blood that can serve as biomarkers for effective vaccine design. We found that DENV sero-positive participants had increased pre-vaccination frequencies of Tfh cells and higher levels of the Tfh related chemokine CXCL13/BLC that plays a role in directing antigen-specific responses. This pre-vaccination Tfh profile and CXCL13/BLC are then correlated positively with the vaccinees’ ability to produce neutralizing antibody against all four sero-types (breadth of the Response) of DENV, an important goal for all DENV vaccine trials.

## Introduction

Dengue virus (DENV), a mosquito borne flavivirus has been considered a worldwide health threat since up to 350 million individuals live in high-risk transmission areas. The rate of Dengue infection has been estimated to be more than 390 million cases annually in tropical and subtropical countries, of which 96 million manifest clinical pathologies (1, 2). Besides the health problem, DENV is a global economic burden (3). In 2013, the total annual cost of symptomatic Dengue cases was approximately $1.51 billion. This economic burden has risen substantially over the years, as the total annual cost of symptomatic Dengue cases in 2016 reached $5.71 billion (4). Billions of lives and dollars could be saved globally by preventative interventions, particularly the development of the Dengue vaccine. Thus, there is an urgent demand for an efficacious Dengue vaccine, and this is illustrated by the international collaborative efforts from various world health organizations and federal institutions.

There are several challenges that hampered the development of a Dengue vaccine. One of which is the presence of 4 different serotypes of DENV, with each having the ability to cause disease. Moreover, the presence of 4 different serotypes causes cross-reactive antibody interaction (5, 6). An effective vaccine must be able to generate neutralizing antibodies against all serotypes of DENV to ensure full protection. Antibody-dependent enhancement can be induced due to pre-existing antibodies in the second infection with a heterogeneous type of DENV (5, 6). There are several vaccine candidates with different platforms under clinical development including live attenuated, purified inactivated, and DNA vaccine platforms. Thus far, only live attenuated candidates had entered phase III trials (7, 8). CYD-TDV, developed by Sanofi Pasteur with the trade name of DENGVAXIA^TM^, is the only vaccine that has completed phase III trials and licensed in many countries worldwide since 2015 (9, 10). CYD-TDV is a tetravalent, live attenuated, recombinant vaccine with a Yellow fever 17D backbone (11). The prM and E proteins of the Yellow fever virus are replaced by the prM and E proteins of 4 serotypes of DENV combined with the nonstructural genes of yellow fever 17D vaccine strain (chimeric yellow fever Dengue – CYD) (12). CYD-TDV has been licensed to be given only to Dengue seropositive individuals due to the observation of disease enhancement among seronegative vaccinated individuals (13). Therefore, Dengue serostatus effect must be further investigated for optimal Dengue vaccine design and development.

T follicular helper cells (Tfh cells) are specialized CD4+ T cells that support the Germinal center (GC) formation and influence B-cell responses (14, 15). The role of Tfh has become the interest of many studies, including vaccination, due to their fundamental role in facilitating the generation of pathogen-specific and long-lasting antibodies (16). Tfh cells enhance GC B-cells function through secreting IL-21 and up-regulating numerous proteins and transcriptional factors such as ICOS, CD40, Bcl-6, and PD-1 (17). The primary location of Tfh cells is within the germinal centers. However, due to lack of non-invasive means to access this compartment, a surrogate of these cells that share similar phenotypic and functional characteristics found in the peripheral blood called circulating Tfh cells (cTfh) are intensively studied (16, 18). The cTfh cells are characterized by the expression of CXCR5, inducible co-stimulator (ICOS) and programmed death-1 (PD-1). The cTfh cells can be evaluated to predict GC’s activity since they have been positively correlated with neutralizing antibodies and vaccine responses (19, 20). cTfh cells are comprised of different subsets; cTfh1, cTfh2, and cTfh17. Each subtype expresses different cytokines and provides distinctive help for B cells (21). All cTfh subsets are capable of providing B cell help with varying degree of efficacy, where cTfh17>cTfh2>cTfh1(22). Thus, cTfh cells provide an excellent tool for monitoring vaccine responses (23).

CXCL13/BLC, also known as B lymphocyte chemoattractant (BLC), is a potent chemokine that selectively chemoattracts B cells. It is expressed by follicular dendritic cells, stromal cells in lymphoid organs, and in GC Tfh cells. CXCL13/BLC recruits cells expressing CXCR5 on their surface (24). Targeting either CXCR5 or CXCL13/BLC results in defects in the formation of the germinal center in lymphoid organs. Therefore, CXCL13/BLC is presumed to direct both B cells and antigen-specific T cells into germinal centers in lymphoid organs. Germinal centers specific Follicular helper T cells (GC-TFH) and B cells express CXCR5 on their surface, a receptor crucial for their migration to the B-cell follicle (25). A strong and direct correlation between plasma CXCL13/BLC and lymphoid GC-Tfh cells has been established previously (26). Additionally, GC-Tfh cells are robust producers of CXCL13/BLC (26). Follicular Helper T cells preferentially secretes CXCL13/BLC to attract B cells to the germinal centers. Various studies on HIV infection revealed the role of CXCL13/BLC as a plasma biomarker of germinal center activity in which elevated plasma levels of CXCL13/BLC associated with the generation of broadly neutralizing antibodies (26, 27). Recently, it has been shown that CXCL13 levels in plasma provides strong indication for germinal centers activities in response to vaccines and infections (28). Thus, measuring CXCL13 in plasma provides critical information on the immune activities occurring in lymphoid tissues.

To investigate the influence of Dengue virus serostatus on vaccine outcome, we have utilized samples from Brazil that were obtained longitudinally during Live attenuated TV003 phase II clinical trial. This cohort is a unique cohort because it could be utilized as a tool to characterize the pre-vaccination basal microenvironment and how it differs between Dengue seropositive and seronegative groups. We investigated both the cellular differences and the immune signatures differences at baseline/pre-vaccination (Day 0) prior to TV003 vaccination in the two groups. The vaccine response in this study was measured by two outcomes, which are the neutralizing antibody titers as well as the breadth of the vaccine response (seroconversion).

## Results

### Variation of the vaccine response among the subjects

To investigate the magnitude of neutralizing antibody response induced by TV003 against the four DENV serotypes, we measured levels of neutralizing antibodies induced by the vaccine via a plaque reduction neutralization test (PRNT) for the whole Phase II cohort. Individuals were dichotomized as Immune (seropositive) and Naive (seronegative) based on their respective capacity to neutralize any of the 4 DENV serotypes prior to vaccination or absence of neutralization. We investigated the neutralizing antibody titers at day 28, 56, and 91 against all four Dengue virus serotypes, which were then integrated using area under the curve pre boost (AUCp) to account to represent the magnitude of the serotype-specific responses. The heatmap visually represents neutralizing antibody titers area under the curve (AUCp) for 266 subjects with vaccine group types and serostatus groups (Fig 1A). The placebo vaccine group is labeled green while the vaccinated group is labeled red. Among the serostatus groups, seropositive group (immune) is labeled light blue while seronegative group (Naïve) is labeled magenta. The neutralizing antibody titers area under the curve (AUCp) is labeled blue, with range of low to high fold change. Qualitative visual representation suggest that seropositive individuals who received the vaccine showed more robust neutralizing antibody titers against all four serotypes compared to their Naïve counterparts, or individuals who received the placebo. Interestingly, we observed large variability in the response amongst all vaccinated subjects suggesting that pre-vaccination environment and conditions could play a role in the response (Fig 1A). To further understand these variations, we used Principal Component Analysis (PCA) that clusters samples in groups based on their similarity (Fig 1B). As expected, placebo controls clustered to the left with minimal neutralizing antibody titers; whereas there is a clear distribution difference between naïve vaccinated (seronegative at pre-vaccination) and Dengue immune (seropositive at pre-vaccination) vaccinated subjects, with most of the higher response observed in the Dengue immune vaccinated subjects. In addition, we observed minor similarities between a) Dengue serotype 2 (DENV2) and serotype 3 (DENV3) specific responses and b) Dengue serotype 1 (DENV1) and serotype 4 (DENV4) (Fig 1B).

**Fig 1:**
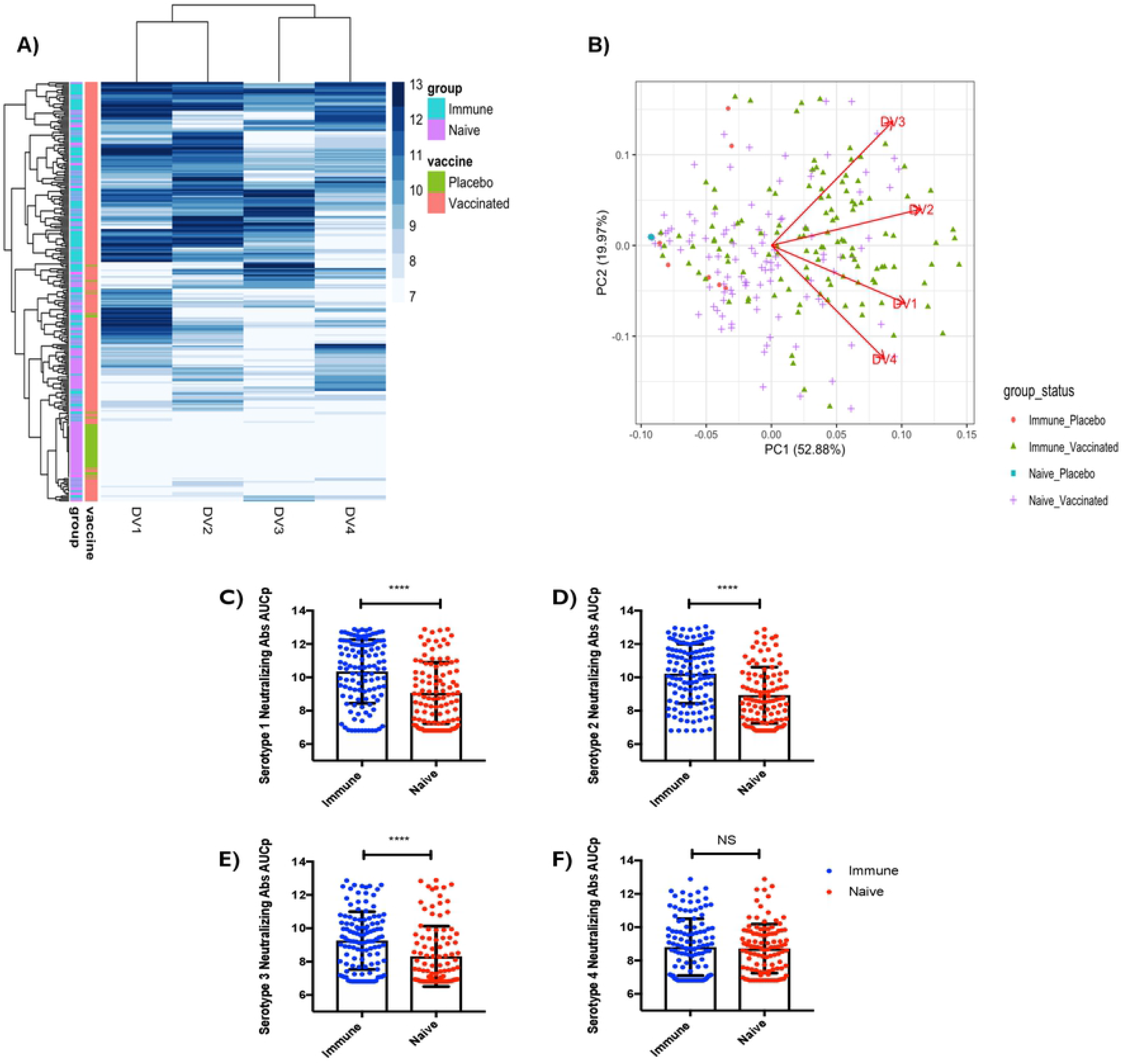
Response to dengue vaccine is variable and impacted by the pre-vaccination immune status. **(A)** Heatmap of subject AUC values. For each subject there are four AUC values represented in DV1, DV2,DV3 and DV4 columns. The higher AUC value is represented by a darker blue strike. Two columns to the left of the graph help to distinguish the vaccine status and which group subjects belong to. **(B)** Biplot for 266 subjects DV1-4 AUC values plotted in two dimensions on first two principal components and DV1-4 vectors indicating each’s contribution to PC1 and PC2. Samples are colored according to their baseline/pre-vaccination immune status and whether they received the dengue vaccine. **(C)** Y axis is Dengue Serotype 1 neutralizing antibody titers (AUCp). **(D)** Y axis is Dengue Serotype 2 neutralizing antibody titers (AUCp). **(E)** Y axis is Dengue Serotype 3 neutralizing antibody titers (AUCp) **(F)** Y axis is Dengue Serotype 4 neutralizing antibody titers (AUCp). This was accomplished using plaque reduction neutralization test (PRNT). (Immune n=119, Naive n=105). Unpaired non-parametric Mann Whitney test, 95% confidence was calculated for **(C-F).** Before vaccination, seropositive subjects with existing antibodies to any of the four Dengue virus serotypes due to prior infection were referred to as (“Immune”), or seronegative subjects with no existing antibodies to any of the four Dengue virus serotypes were referred to as (“Naïve”).

We further, demonstrate quantifiably that pre-existing dengue infection of any of the four serotypes before vaccination was associated with significant increase in neutralizing antibodies against serotypes DENV1, DENV2, and DENV3 post vaccination, indicating an increased ability for previously Dengue seropositive (Immune) individuals to influence neutralizing antibody titers post vaccination. As shown in (Fig 1 C, D, E), presence of Dengue neutralizing antibody titers to any of the four serotypes at the baseline/pre-vaccination time point result with significant increase in DENV1 (p<0.0001), DENV2 (p<0.0001), and DENV3 (p<0.0001) when compared to individuals who are not positive for any of the Dengue serotypes. Notably, the presence of antibodies against any of the four serotypes at baseline/pre-vaccination time point did not correspond to the induction of significant neutralizing antibody titers against DENV4 post vaccination (Fig 1F). This could be likely due to the fact that this cohort has much less exposure to DENV4 due to the prevalence of of other serotypes circulating this particular geographical locations. From this data, we can infer that immunization with this vaccine is capable of eliciting robust immune responses in seropositive individuals against majority of the serotypes. Overall, using unsupervised analysis, we observed qualitatively that seropositive (Immune) individuals seem to elicit more vaccine induced neutralizing antibody titers as compared to their seronegative (Naïve) counterparts (Fig 1 A, B). Therefore, we next investigated the immune and naïve neutralizing response AUCp quantitatively and found that pre-existing Dengue virus seropositivity is associated with significant increases in neutralizing antibodies against DENV1, DENV2, and DENV3 post-vaccination (AUCp) (Fig 1C-F).

### Seropositive (Immune) individuals exhibited higher levels of serotype-specific neutralizing antibody titers (AUCp) after vaccination compared to seronegative (Naïve) individuals

To help uncover how seropositivity affected post vaccine antibody outcome, we further stratified the seropositive subjects (Immune) based on presence of individual Dengue subtypes specific titers prior to vaccination. Our results show that pre-existing antibody titers against Dengue virus is associated with significant increases in neutralizing antibody responses in a serotype-specific manner for three of the four serotypes. Individuals who were seropositive for DENV1 before vaccination, showed significantly higher levels of DENV1-specific nAbs (p<0.0001) and DENV3-specific nAbs (p=0.0436) post vaccination when compared to seronegative individuals (Fig 2A, 2C). Interestingly, we did not observe any significant increase in DENV2 or DENV4-specific nAbs in those subjects post vaccination (Fig 2B, 2D). Additionally, individuals who were seropositive for DENV3 before vaccination, showed significantly (p<0.0001) higher levels of DENV3 specific nAb (Fig 2G) and no difference in DENV-1,2,4-specific nAbs post vaccination. Similarly, individuals who were seropositive for DENV4 before vaccination, showed significantly (p=0.0061) higher levels of DENV4-specific nAbs (Fig 2L). However, we did not observe any increase in DENV1, 2, 3-specific nAbs in those subjects (Fig 2 I-K). This serotype-dependent response was not seen in individuals who were previously immune to DENV2 (Fig S1). We suspect that this is due to the fact that there were fewer subjects in the seropositive cohort that are seropositive only to DENV2.

**Fig 2:**
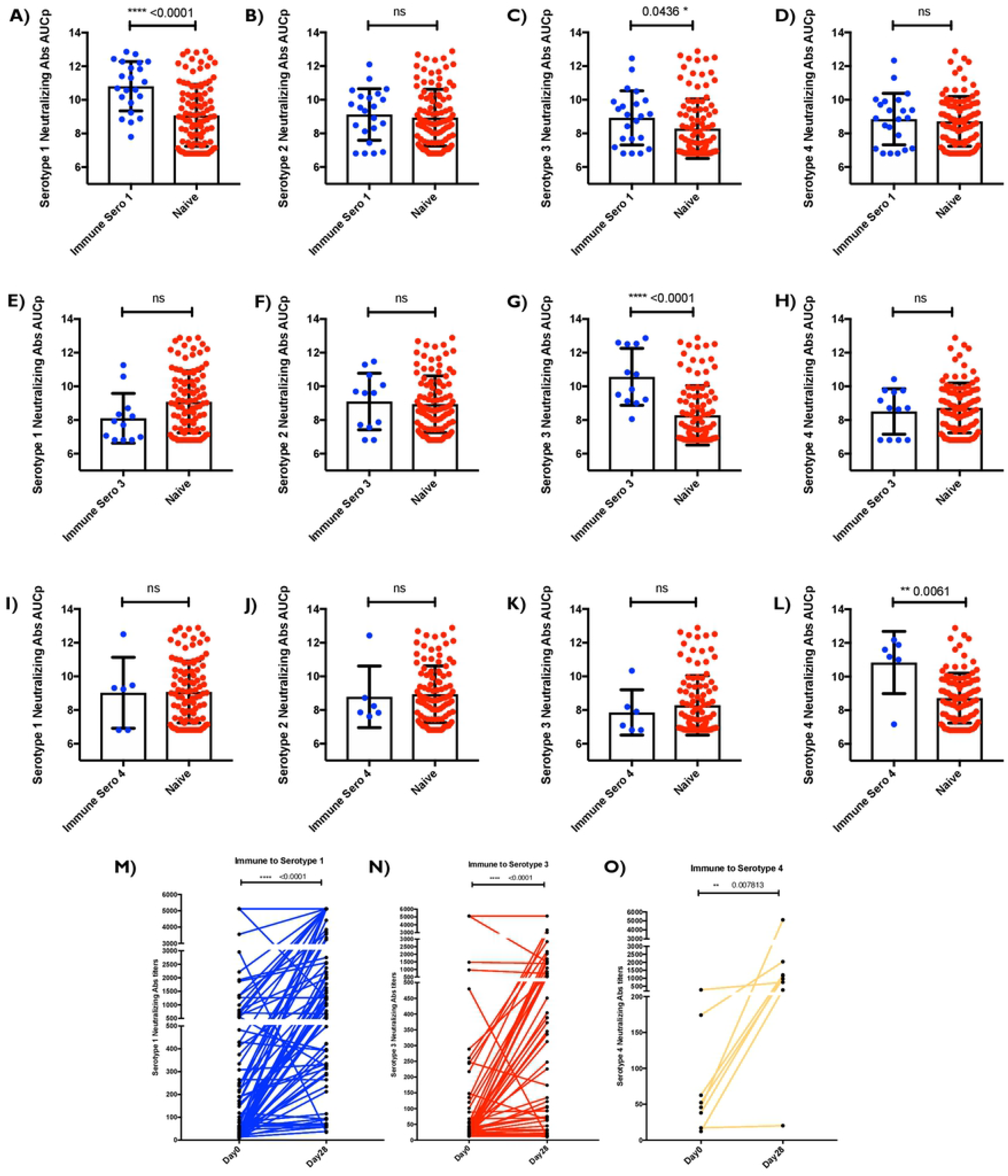
TV003 vaccine induced neutralizing antibody titers AUCp is dependent on pre-vaccination serotype. **(A, B, C &D)** “Immune Sero 1” are subjects who only had Dengue Serotype I neutralizing antibody titers (AUCp) at pre-vaccination/baseline time point. **(E, F, G &H)** “Immune Sero 3” are subjects who only had Dengue Serotype 3 neutralizing antibody titers (AUCp) at pre-vaccination/baseline time point. **(I, J, K &L)** “Immune Sero 4” are subjects who only had Dengue Serotype 4 neutralizing antibody titers (AUCp) at pre-vaccination/baseline time point. Seronegative subjects with no existing antibodies to any of the four Dengue virus serotypes at pre-vaccination/baseline time point are referred to as “Naïve”. **(A,E &I)** y axes are Dengue Serotype I neutralizing antibody titers (AUCp). (B,F &J) y axes are Dengue Serotype 2 neutralizing antibody titers (AUCp). **(C,G &K)** y axes are Dengue Serotype 3 neutralizing antibody titers (AUCp) **(D,H &L)** y axes are Dengue Serotype 4 neutralizing antibody titers (AUCp). **(M)** Neutralizing antibody titers at day 0 and day 28 for dengue serotype 1. **(N)** Neutralizing antibody titers at day 0 and day 28 for dengue serotype 3. **(O)** Neutralizing antibody titers at day 0 and day 28 for dengue serotype 4. This was accomplished using plaque reduction neutralization test (PRNT). Unpaired non-parametric Mann Whitney test 95% confidence was calculated for **(A-L).** Paired non-parametric Wilcoxon test, 95% confidence was calculated for **(M-O).**

Neutralizing antibody titers measured on Day 0 pre-vaccination and Day 28-post vaccination show that there is a clear and significant difference between pre-vaccination antibody levels and vaccine-induced antibody levels (P<0.0001, P<0.0001, P=0.0078). This suggests that the increased antibody titers were in fact, the result of immunization rather than pre-existing levels of nAb at pre-vaccination (Fig 2 M, N, O).

Together, this data suggests that this vaccine candidate triggers a robust serotype specific immune response in which seropositive individuals generate significantly high neutralizing antibody titers against the serotype that they were previously exposed to.

### Higher baseline/pre-vaccination frequency of total cTfh in the seropositive (Immune) than seronegative (Naïve) group

To better understand the immunological components that might be influencing vaccine outcome as result of sero-status, we classified our cohorts into “Immune” (presence of neutralizing antibody titers to any of the four serotypes at baseline/pre-vaccination time point) or Naive (absence of neutralizing antibody titers to any of the four serotypes at baseline/pre-vaccination time point). Given the complexity of immunophenotyping and the paucity in samples from all vaccinees, we carried out FACS based analyses in a sub-set of the cohort (n=44 Naïve and n=14 Immune) at the baseline/pre-vaccination. First, we investigated the T cell compartment, and found no significant differences in any of the major T-cell subsets such as total CD4+ T cells, total memory cells, total CD3+CD4- cells that differentiate Immune and Naïve subjects (S3 Fig). However, we detected a significantly higher (p=0.0082) frequency of total circulatory Tfh cells (cTfh; marked by CD3+CD4+CD45RA-CXCR5+ expression) in the subjects that belonged to the Immune group compared to those in the Naïve group (Fig 3A). While not statistically significant, it is intriguing to note that subtypes of cTfh cells were different in Immune vs Naïve subjects. One of the subtypes of cTfh cells, cTfh1 (marked by expression of CXCR3 within the cTfh population) seems to be trending higher in the Immune group, while cTfh2 (marked by absence of CXCR3/CCR6 expression within the cTfh population) seems to be trending higher in the Naïve group (p= 0.0786, p=0.0505) (Fig 3 B and C). This data suggests that this basal difference of the total cTfh and cTfh sub-populations could be potential immunological underpinnings that might explain serostatus mediated differences in vaccine responses. Based on our and others previous reporting, total cTfh cells have the ability to provide B cell help using in vitro co-culture assays (18, 23, 29). Therefore, we pursued this subset further in this study.

**Fig 3:**
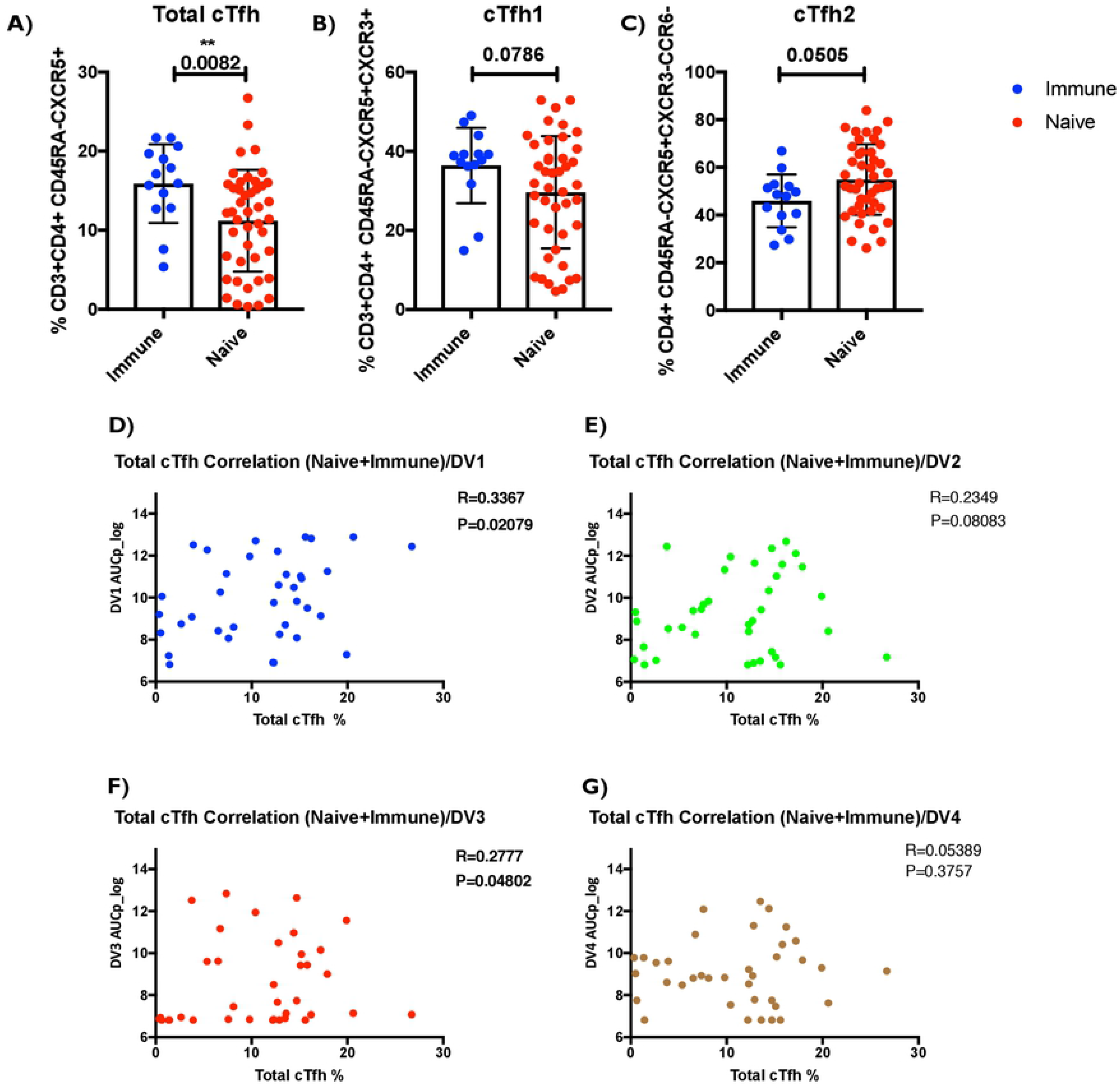
Higher baseline/pre-vaccination frequency of total cTfh in the seropositive (Immune) than seronegative (Naïve) group, positively correlate with neutralizing antibody titers AUCp. **(A)** Y axis is the total cTfh Frequency CD3+ CD4+ CD45RA- CXCR5+. **(B)** Y axis is the cTfh1 frequency CD3+ CD4+ CD45RA- CXCR5+ CXCR3+ CCR6-. **(C)** Y axis is the cTfh2 frequency CD3+ CD4+ CD45RA- CXCR5+ CXCR3- CCR6-. **(D)** Total cTfh frequency (CD3+ CD4+ CD45RA- CXCR5+) correlation with dengue serotype 1 AUCp. **(E)** Total cTfh frequency (CD3+ CD4+ CD45RA- CXCR5+) correlation with dengue serotype 2 AUCp. **(F)** Total cTfh frequency (CD3+ CD4+ CD45RA- CXCR5+) correlation with dengue serotype 3 AUCp. **(G)** Total cTfh frequency (CD3+ CD4+ CD45RA- CXCR5+) correlation with dengue serotype 4 AUCp. Pre-vaccination/baseline subjects PBMCs and optimized T cell panel for ex-vivo staining was utilized. 1 million cells per panel were stained. (Naïve n=44, Immune n=14.) Unpaired non-parametric Mann Whitney test, 95% confidence was calculated for **(A-C).** Non-parametric Spearman Correlation (R=rho) with 95% confidence Interval was calculated between Total cTfh frequencies and Area Under the Curve (AUCp) for DV serotypes 1-4. (n=37 for **(D-G))**. Before vaccination, seropositive subjects with existing antibodies to any of the four Dengue virus serotypes due to prior infection were referred to as (“Immune”) or seronegative subjects with no existing antibodies to any of the four Dengue virus serotypes were referred to as (“Naïve”).

### Pre-vaccination frequency of total cTfh positively correlates with neutralizing antibody titers (AUCp) and the breadth of the vaccine response

Given the known role of cTfh in mediating B cell immunity and the increased propensity of seropositive patients to generate a higher magnitude of serotype-specific responses, we investigated if cTfh frequencies were contributing to that effect. We investigated whether there is correlation between total cTfh (marked by CD3+CD4+CD45RA-CXCR5+ expression) frequency at baseline/pre-vaccination and neutralizing antibody responses at post vaccination time points. We found that the cTfh frequency positively correlated with neutralizing antibody titers area under the curve AUCp (Fig 3). Specifically, we found that total cTfh frequency significantly and positively correlated with AUCp for DENV1 (P<0.05) and DENV3 (P<0.05) (Fig 3D and F) indicating that having a higher frequency of total cTfh at pre-vaccination might predict higher Dengue neutralizing antibody titers elicited via the TV003 vaccination. Correlation of cTfh frequency and DENV2 and DENV4 specific neutralizing antibody titers showed positive correlation but did not result in statistical significance (Fig 3E and G).

We next determined whether frequency of cTfh cells at pre-vaccination correlates with breadth of the vaccine response (defined by the vaccinees’ ability to mount high neutralizing antibody titers against all four DENV serotypes). First, we profiled for CD4+ T cell subsets using tSNE based FACS data analysis we found 23 distinctive clusters depicted in (Fig 4A). The 23 distinctive clusters are displayed as a heatmap in (Fig 4B) that included cluster 16 that represents circulating Tfh cluster (Fig 4B). We further investigated T-cell subsets/status at baseline/pre-vaccination time point and correlated with the breadth of Dengue virus specific vaccine responses. This was done by separating the baseline/pre-vaccination frequencies of cTfh between subjects who mounted neutralizing antibodies against all four serotypes (termed “high” breadth as defined previously) compared to subjects who mounted neutralizing antibodies against <3 serotypes (termed “low” breath as defined previously). Uniquely, circulating Tfh cells robustly correlated with the breadth of Dengue virus specific neutralizing antibodies induced upon vaccination as represented by cluster 16. This analysis confirmed that cTfh frequencies are statistically significantly (p<0.05) higher in “high” subjects compared to “low” subjects suggesting this cell type’s important role in the breadth of vaccine specific immune responses (Fig 4C).

**Fig 4:**
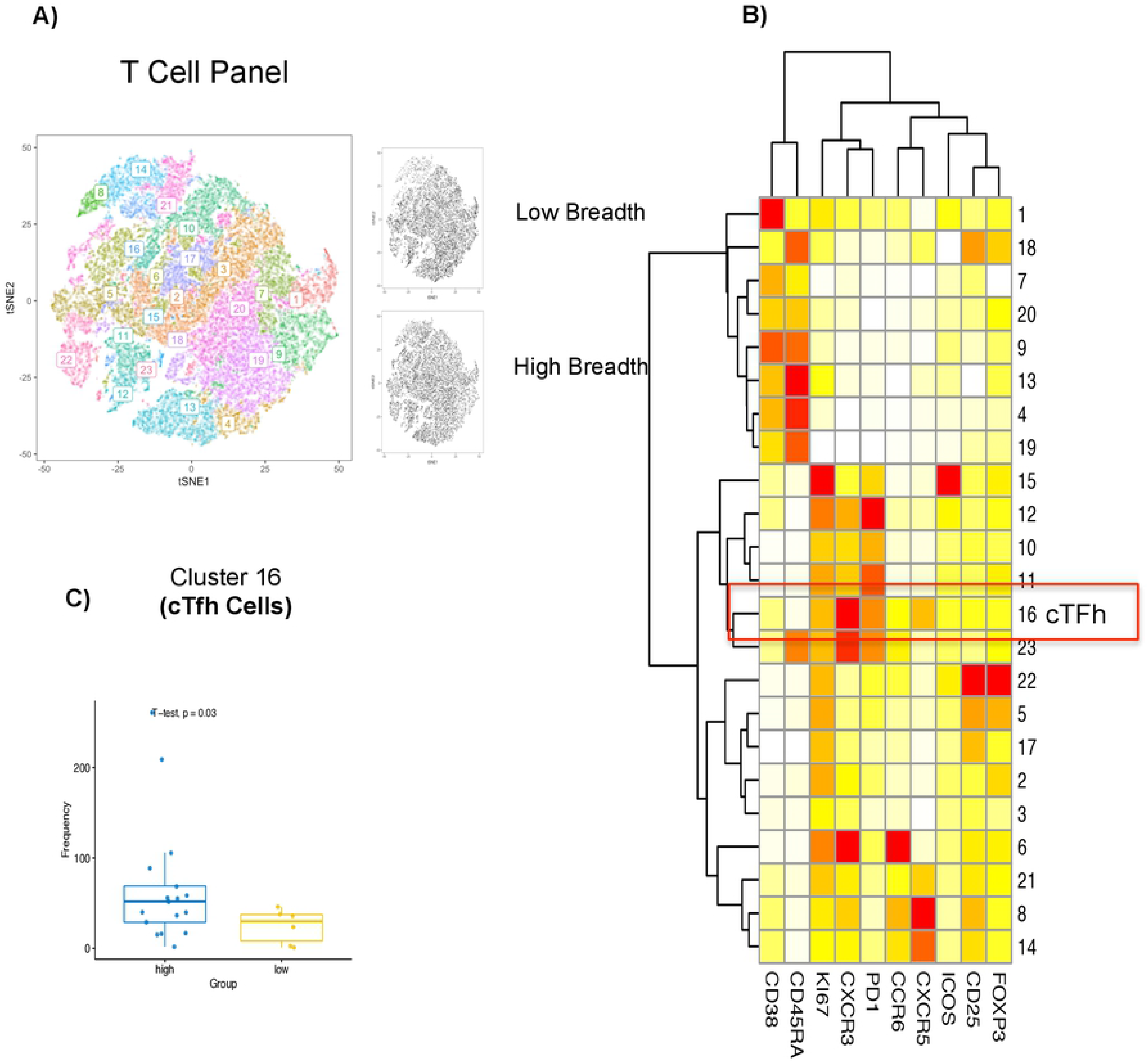
Pre-vaccination frequency of total cTfh positively correlates with the breadth of the vaccine response. **(A)** Cell cluster distribution depicts 23 separate distinctive clusters**. (B)** Heatmap of the 23 different clusters expressing different phenotypes using the optimized T cell panel with markers denoted in the X-axis. Cluster 16 represents a cTfh cluster. **(C)** Boxplot of the significant cluster (cluster 16) investigating the cluster with the breadth of the vaccine response. Subjects who mounted neutralizing antibodies against all four serotypes (termed “high” breadth) were compared to subjects who mounted neutralizing antibodies against < 3 serotypes (termed “low” breadth). Cells were gated on live CD4+CD3+ cells and the expression was scaled and used to perform clustering and was visualized using t-Distributed Stochastic Neighbor Embedding (tSNE). Cell frequency per cluster/ per sample was derived and used for statistical testing between high and low breadth across donors. Day 0 donor samples were associated with breadth of the response post vaccination. Welch’s T-test was performed for the breadth of vaccine response **(C)**.

### CXCL13/BLC expression levels at pre-vaccination are associated with high breadth of vaccine response

After recognizing the positive association between cTfh profile with vaccine response, we next investigated whether secreted immunomodulatory factors such as cytokines and chemokines at baseline/pre-vaccination time point could also provide information on pre-vaccine environment. Therefore, we used a multiplexed approach and profiled 65 cytokines and chemokines to examine correlation with vaccine response. Among all these analytes evaluated, B lymphocyte chemoattractant (BLC) or CXCL13 (secreted by multiple cells including Tfh cells) showed a difference in detection between immune and naïve subjects (Fig 5A) with statistical significance of p<0.005. Interestingly, it has previously been shown that CXCL13/BLC or BLC is a plasma biomarker of GC activity and that it is associated with the generation of broadly neutralizing antibodies against HIV in HIV-infected individuals (26). As GC are required for almost all B cell receptor affinity maturation and the optimization of B cell antibody responses, this is a critical parameter to monitor for vaccine responses.

**Fig 5:**
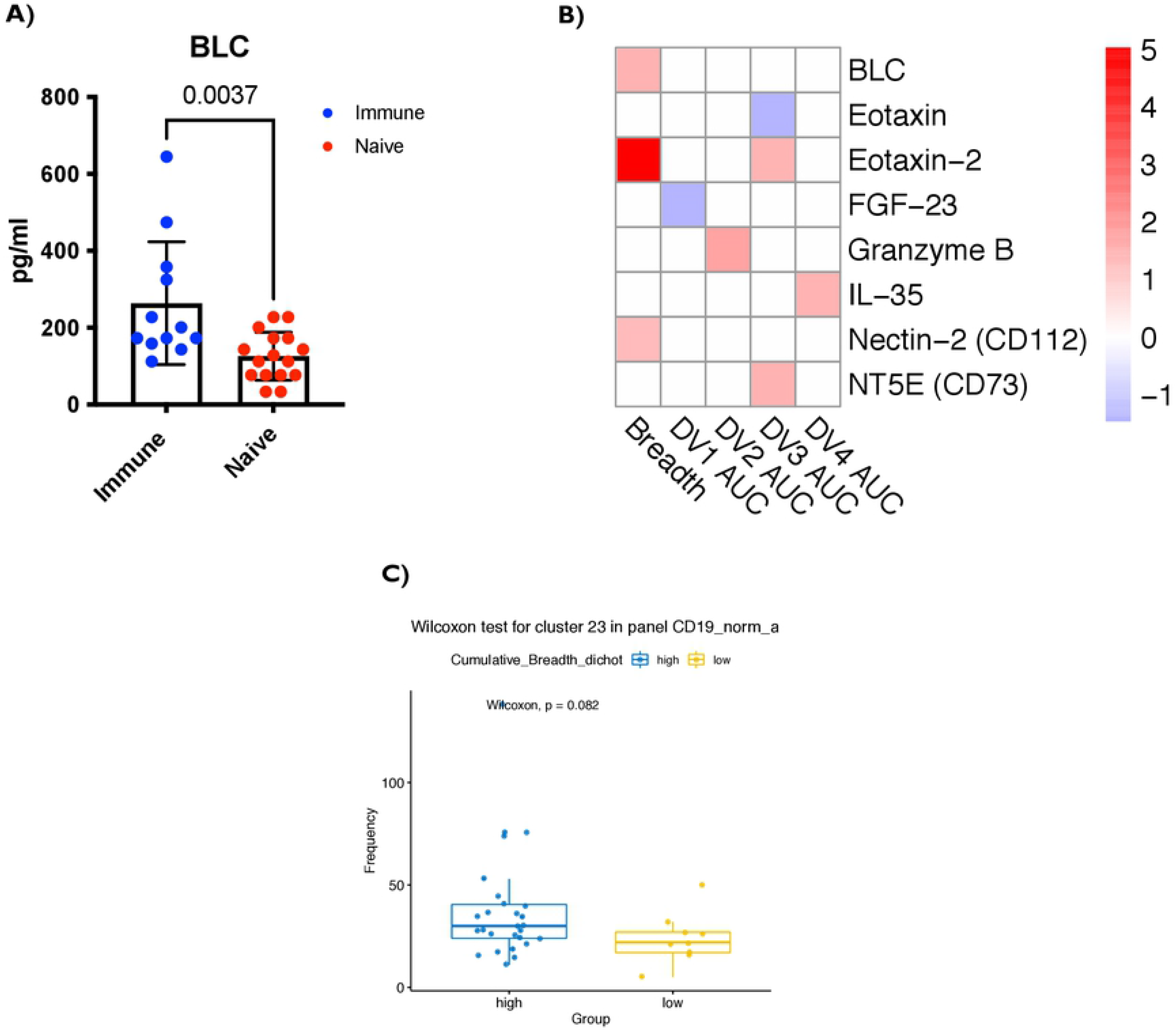
Higher CXCL13/BLC expression levels at pre-vaccination are associated with the breadth of the vaccine response. **(A)** Y axis shows CXCL13**/**BLC levels at pre-vaccination/baseline in pg/ml. Unpaired non-parametric Mann Whitney test, 95% confidence was calculated. **(B)** Heatmap of cytokines and chemokines are shown that are significantly correlated with neutralizing antibody titres (AUCp) as well as the breadth. The x axis of the heatmap has the neutralizing antibody titer (AUCp) for the four Dengue serotypes as well as the breadth. To the right of the heat map is the color gradient Signed log10 p-values (−log10 (p’) x sign of the Welch’s T-test or rho coefficient, respectively), which was used to show the strength of the association to each outcome. For the analytes in (**A and B)**, neat Plasma samples were used. (Immune n=12, Naïve n=16.) **(C)** Representation of cluster 23 (Plasmablast) pre-vaccination/baseline subject samples were associated with breadth of the response post vaccination. This is cluster 23 of the t-Distributed Stochastic Neighbor Embedding (tSNE) for B cell panel in supplementary figure 4. Pre-vaccination/baseline subjects PBMCs and optimized B cell panel for ex-vivo staining was utilized. 1 million cells per panel were stained. (Naïve n=44, Immune n=14). Before vaccination, seropositive subjects with existing antibodies to any of the four Dengue virus serotypes due to prior infection before were referred to as (“Immune”) or seronegative subjects with no existing antibodies to any of the four Dengue virus serotypes were referred to as (“Naïve”). Paired non-parametric Wilcoxon test, 95% confidence was calculated for **(C).**

We further performed an in-depth characterization of cytokines and chemokines that may significantly correlate with antibody titers at various time points post vaccination (Fig 5B). We next evaluated which cytokines/chemokines at baseline time point correlated with the breadth of responses at various time points post vaccination. Remarkably, BLC/CXCL13, showed a statistically significant correlation with the breadth of response. Moreover, Eotaxin-2 and Nectin-2 also showed statistically significant correlation with the breadth of response. We observed few interesting baseline cytokine/chemokine profiles with individual serotype specific neutralizing antibody titers that were statistically significant: i) FGF-23 correlated with AUCp of neutralizing antibody response to DENV1, ii) Granzyme B correlated with AUCp neutralizing antibody responses to DENV2, iii) there were many analytes such as Eotaxin, Eotaxin-2 and NT5E that correlated with AUCp neutralizing antibody responses to DENV3, and iv) IL-35 correlated with AUCp neutralizing antibody responses to DENV4. As stated previously, evoking a higher breadth in response (i.e. neutralizing all four serotypes) is a critical outcome for a successful vaccine. This further emphasizes the critical role that cTfh related immunological factors that might play in influencing vaccine specific immune responses.

### B-cell plasmablast frequency at baseline/pre-vaccination time point correlates with the breadth neutralizing antibody response

In addition to performing T-cell related immunophenotyping, we investigated whether there is correlation between frequency of B-cells (and subset) at baseline/pre-vaccination time point with serostatus and favorable vaccine responses. Evaluation of B-cell frequencies with favorable vaccine responses revealed that B-cell plasmablast (characterized by CD19+CD38+CD27+ expression) frequency showed positive correlation with AUCp for DENV3 and breadth of neutralizing antibodies against all four serotypes (S4D Fig). While not statistically significant, we observed increased trend in plasmablasts in “high” subjects compared to “low” subjects (Fig 5C) (p=0.082). Notably, ee observed that the majority of the B-cell subsets measured, including total CD19+ B-cells (S4A Fig), total class-switched B-cells characterized by CD19+CD10-IgM-IgD-expression (S4B Fig), and activated memory B-cells, characterized by CD19+CD10-IgM-IgD-CD21-CD27+(S4C Fig), did not show any statistically significant difference between immune and naïve subjects.

### Monocytes correlate positively with both the AUCp and the breadth of the response

Innate cells are critical players both in the context of Dengue virus infection and in the context of mounting favorable responses to vaccination. As part of the immunophenotyping panel, we evaluated correlations between various innate immune cell subsets (& their functional status) at the baseline/pre-vaccination with serostatus and vaccine responses. Interestingly, using tSNE based FACS data analysis depicted as a heatmap, we observed multiple clusters of monocytic cell subsets that were statistically significantly altered (Fig 6A). These included: i) classical monocytic cell cluster 5 (CD14+CD16-CD163lo) that correlated with neutralizing antibodies AUCp of DENV2 and detected at increased frequency in immune subjects (p<0.01) as compared to naïve subjects (Fig 6B), ii) classical monocytic cell cluster 13 (CD14+CD16-CD163high) that correlated with neutralizing antibodies AUCp of DENV1 and detected at increased frequency in immune subjects (p<0.001) as compared to naïve subjects (Fig 6C), iii) transitional monocytes cluster 11 (CD14+ CD16+) and non-classical monocytic cell cluster 22 (CD14-CD16+) that correlated with neutralizing antibodies AUCp of DENV4 and with higher breadth of vaccine responses as defined previously(Figs 6D and 6E), and (iv) mDC2 cluster 3 (CD14-CD16-CD11c+) that correlated with neutralizing antibodies AUCp of DENV2 and detected at increased frequency in immune subjects.

**Fig 6:**
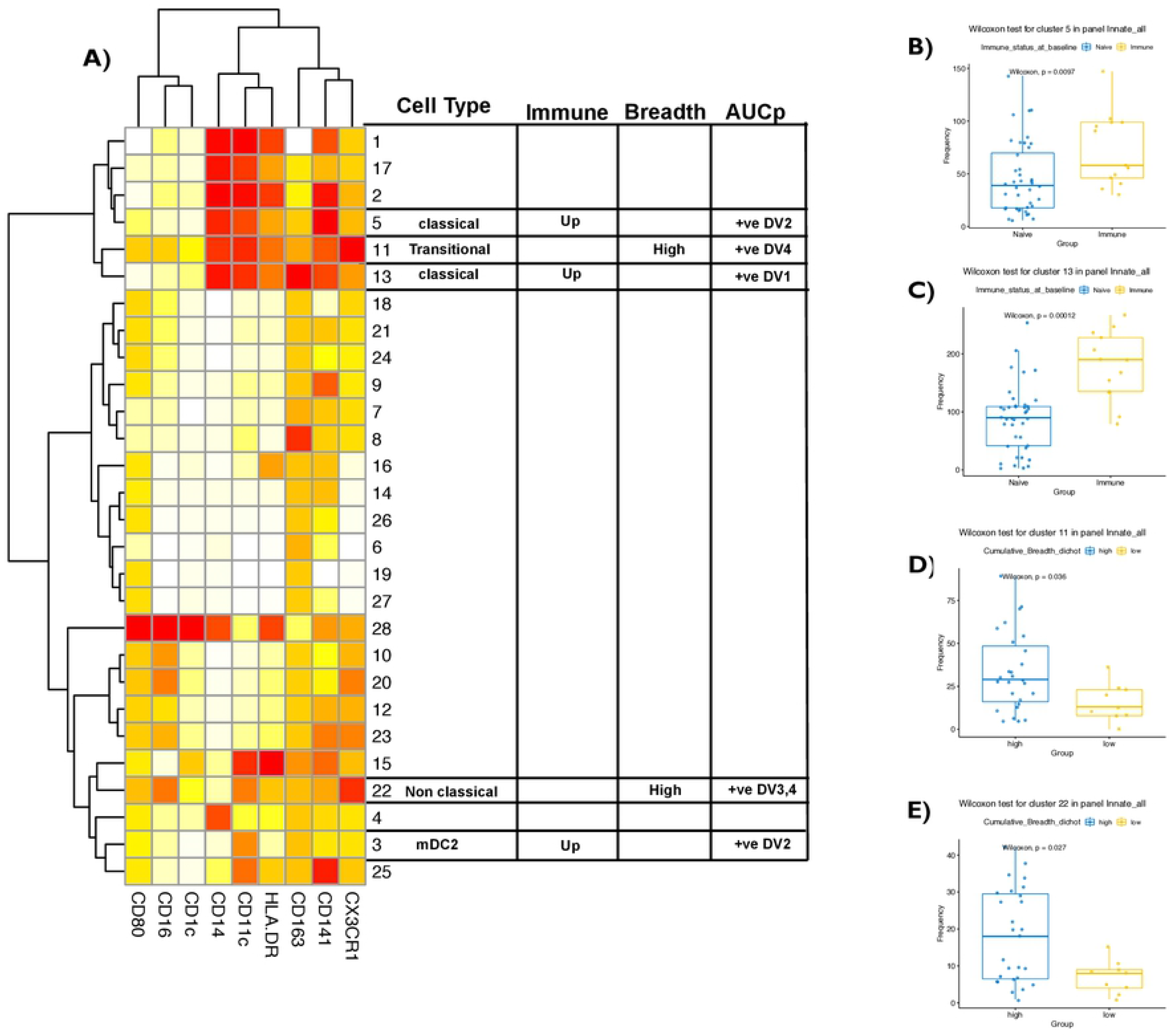
Monocytes correlate positively with both the AUCp and the breadth of the response. **(A)** Heatmap of the 28 different clusters expressing different phenotypes using the optimized innate cell panel are shown with markers in the x axis. Among them are clusters 3, 5,11,13 and 22 that represent type 2 myeloid dendritic cell, classical monocyte, transitional monocyte and non classical monocyte. The table identifies cell type, immune group pre-vaccination/baseline difference, breadth and neutralizing antibody titers (AUCp). Cells were gated on live cells and the expression was scaled and used to perform clustering which was visualized using t-Distributed Stochastic Neighbor Embedding (tSNE). Cell frequency per cluster/ per sample was derived and used for statistical testing between high and low breadth across subjects. Subjects who mounted neutralizing antibodies against all four serotypes (termed “high” breadth) were compared to subjects who mounted neutralizing antibodies against < 3 serotypes (termed “low” breath). Pre-vaccination/baseline subject samples were associated with breadth of the response and neutralizing antibody titers (AUCp) post vaccination. **(B and C)** Frequency of cluster 5 and 13 (classical monocytes) between Immune vs Naïve at pre-vaccination/baseline time point. **(D)** Representation of cluster 11 (transitional monocytes) pre-vaccination/baseline subject samples were associated with breadth of the response post vaccination**. (E)** Representation of cluster 22 (non-classical monocytes) pre-vaccination/baseline subject samples were associated with breadth of the response post vaccination. Before vaccination, seropositive subjects with existing antibodies to any of the four Dengue virus serotypes due to prior infection were referred to as (“Immune”) or seronegative subjects with no existing antibodies to any of the four Dengue virus serotypes were referred to as (“Naïve”). Paired non-parametric Wilcoxon test, 95% confidence was calculated for **(B-E).** Non-parametric Spearman Correlation with 95% confidence intervals was used for correlations with neutralizing antibody titers (AUCp column).

These observations have many important implications based on existing literature on the role of these innate immune populations. Non-classical CD14^dim^ CD16^-^ monocytes are instrumental in the anti-viral response and patrol the vascular endothelium in response to infection or injury and participate in T cell stimulation and proliferation. Classical CD14^+^CD16^-^ monocytes are pro-inflammatory superior phagocytes that have roles in response to stimuli. We believe that this difference in the baseline/pre-vaccination frequency of important anti-viral innate cells like CD14^+^ CD16^-^ monocytes play a role in the downstream recruitment of cTfh and B cells in GC responses.

## Discussion

Among many, a unique challenge facing Dengue virus vaccine development is the phenomenon of antibody dependent enhancement, or ADE. It is documented that pre-existing Dengue virus specific immunity is known to have a much severe adverse outcome in patients that succumb to secondary infection by a different Dengue virus serotype (30). Similarly, the pre-immunization microenvironment of subjects is also known to influence vaccine efficacy, exemplified by observations from Sanofi Pasteur’s CYD-TDV (13). This licensed vaccine has been shown to be effective in preventing severe disease in seropositive vaccinees but could also result in unfavorable outcomes in seronegative individuals (13). As a result, CYD-TDV is only licensed for use in seropositive individuals. However, specific cell types and immune mechanisms that govern the differential outcome in Dengue virus seropositive versus seronegative subjects who receive the vaccine is incompletely understood. In this study, we investigated the effect of Dengue serostatus in a unique Phase II cohort immunized with the TV003 live-attenuated vaccine candidate utilizing both baseline/pre-vaccination and post-vaccination timepoints. Comparison between Immune and Naïve individuals who received either the vaccine or a placebo demonstrated clear differences in AUCp of neutralizing antibodies (nAb), demonstrating the ability of the vaccine to elicit immune responses from immune and naïve subjects, despite a predictable variability between the human donor samples by PCA analysis.

Neutralizing antibody responses were different depending on pre-vaccination serostatus, with seropositive individuals showing demonstrably higher levels of antibody against three of the four-Dengue serotypes after immunization with the vaccine candidate. Pre-existing immunity to one of the four serotypes not only resulted in post-immunization increases of nAbs for that specific serotype. This finding suggests that an immune memory component that synergizes with the TV003 vaccine.

Follicular helper T cells are a crucial part of the adaptive immune response, facilitating the production of high affinity, neutralizing antibodies. As such, their frequency was compared between immune and naïve subjects to better understand how seropositivity affects neutralizing antibody titers. Due to their function in effecting broadly neutralizing antibodies in response to antigenic stimulation, it was surprising to find that seropositive individuals had higher basal frequencies of cTfh cells in circulation than seronegative individuals. It implies that prior dengue infection boosts cTfh homeostasis as a whole, independent of their specificity. The skew in frequency towards cTfh1 cells in the seropositive cohort and towards cTfh2 in the seronegative cohort, while not significant, demonstrated a trend in the seropositive cohort towards an early antiviral immune response, which is crucial for the control of Dengue infection (31). The basal levels of cTfh also correlated with the breadth of the vaccine response and highlights the critical role that cTfh potentially plays in influencing the vaccine specific immune tone.

Interestingly, though B cell help is more efficiently provided by cTfh2 and cTfh17 cells, the cTfh cells that corresponded to high breadth highly expressed the marker CXCR3, which is classically found on cTfh1 cells and not typically considered to be efficient B cell helpers (31). Despite this, similar findings have been made in the study of HIV controllers in which patients demonstrate increased correlations between circulating CXCR3+PD-1^low^ Tfh-like cells and breadth of neutralizing antibodies against HIV-1 (32). Circulating Tfh1 cells have also been implicated in the vaccine-induced immune response against yellow fever (33). Further studies will require a greater elucidation of cTfh subsets and their roles in neutralizing antibody generation in naïve and immunized individuals. Ultimately, the overall positive correlation between cTfh cells and neutralizing antibody levels strengthens the argument that cTfh cells modulate the serotype effect. These findings demonstrate that the frequency of total cTfh cells in subjects’ PBMCs is predictive of vaccine outcome and correlates with a level of protection against all four Dengue serotypes when elicited by the vaccine. It is important to note that while total cTfh frequency and function were identified in this study as correlates to breadth and neutralizing antibody responses, moving forward it will be prudent to assess the function of antigen-specific Tfh cells in response to Dengue peptides at various time points prior to and post-immunization with TV003.

The association between Tfh cells and vaccine specific immune responses is further strengthened by our finding that the key Tfh chemokine, CXCL13/BLC is also positively correlated with the breadth of the immune response. CXCL13/BLC has been previously shown to be an effective plasma biomarker for increased germinal center (GC) activity in mice, non-human primates, and humans in immunization and infection models (26, 34–36). Beyond the association with neutralizing antibody titers, the correlation with the breadth, or the seroconversion to all four serotypes is a significant finding and has meaningful implications in the development of effective Dengue vaccines.

Plasmablasts represent a critical subset of B-cells that actively secrete antibodies in response to vaccination. One key exploration in our study is to evaluate if baseline/pre-vaccination time point of B-cell subsets and more specifically plasmablasts are differentially abundant between Immune and Naïve subjects. Although we observed a trend in plasmablasts being higher in immune subjects, we do not see a statistically significant trend in this cell population. This highlights the fact that the presence of pre-existing memory B-cells specific to DENV (not total memory B-cell frequencies) is more likely an influencer of the difference in neutralizing antibodies induced post vaccination rather than the mere presence of different B-cell subsets. However, it is likely that plasmablasts were significantly altered at post vaccination time points; unfortunately, due to study limitations and lack of the desired sample availability, we were unable to interrogate this question at this moment.

The role of innate immune cells, specifically, monocytes in the Tfh microenvironment and in Dengue vaccine responses needs to be further investigated. The positive correlation between non-classical and transitional/intermediate monocytes and breadth of the immune response suggests a role for these cells in the same pathway in which the cTfh cells participate. Research in mice has demonstrated that the sheer number of IL-12 producing inflammatory monocytes in draining lymph nodes makes them the most dominant IL-12 producers in the dLNs and that T cells reorganize preferentially within the microenvironment toward these signals (37). Research in humans has shown that non-classical monocytes, which are mobile in nature and patrol the endothelium in search of antigen, also have pro-inflammatory behavior and participate in antigen-presentation and T cell stimulation (38–40). This ability also extends to intermediate monocytes that have also been shown to stimulate T cell proliferation (41–44).

While our entire work is in Dengue virus infected/vaccinated subjects, we strongly believe that there are numerous implications for our findings in the context of SARS-CoV-2 vaccine development. Due to the known homology between the RBD of SARS-CoV-2 and SARS-CoV-1, there is potential for ADE to occur. A study recently has demonstrated that ADE occurs through neutralizing antibodies against RBD (45, 46). Therefore, it is worth investigating if the significant adverse events (including mortality) in some populations could be associated with potential pre-exposure to one or more milder strains of coronaviruses, including common-cold coronaviruses (47).

Though questions remain about the actions of innate immune cells including classical, non-classical, and transitional/intermediate monocytes, these factors are shown to have a relationship with the cTfh cell frequency, neutralizing antibody titers, and breadth of the immune response. The importance of innate immune cells in potentiating a pro-Tfh microenvironment is clear, and we predict that the role of circulating monocytes is only just coming to light. From these data, we conclude that monocytes play a key role in the Tfh microenvironment. This unique interplay between monocytes and cTfh cells will likely enable efficient B cell help within the GCs that ultimately results in the persistence of memory B and plasma cells (and a corresponding increase in high affinity, neutralizing antibodies). With seropositive subjects demonstrating increased baseline/pre-vaccination cTfh levels and neutralizing antibody titers, we have shown that the serostatus effect is likely the result of the discrepancies in cTfh levels between Immune and Naïve subjects. Together, these studies will help delineate the mechanisms by which serostatus affects immune responses to dengue vaccine and how our unique signature of cTfh and CXCL13/BLC levels at baseline may be predictive of vaccine outcomes.

Prediction of vaccine outcomes would lead to identification of targets that could improve and predict vaccination responses, which would be necessary in heterogenic response typically observed in a population. There are many studies that investigate baseline signatures to help predicting favorable vaccine outcome in different diseases models. In influenza for example, this approach has help to look at preexisting antibody titers and variation in response to vaccine outcome (48), and immune cell subpopulation phenotypes in vaccine outcomes (49). In the context of subjects vaccinated with a hepatitis B virus vaccine, specific gene transcription profiles and pro-inflammatory status at the time of vaccination were correlative of poor vaccine outcome (50). In SIV models, a cytokine signature was uncovered as predictive of vaccine efficacy and outcome (51). Assessment of immune sub-population signatures in mRNA vaccine against COVID response showed a monocyte-specific role that correlated to neutralizing antibodies, while inflammatory response-based gene transcription correlated as a vaccine signature that overlapped with responses elicited by other vaccines (52), giving a possible baseline candidate that could be used for predicting vaccine outcomes in multiple models. In this study, we have identified cTfh CXCL13/BLC, and Innate cells as important signatures that correlate to neutralizing antibody breadth and thus a candidate to predict vaccine outcome.

Finally, we propose a unique mechanistic working model that may provide context regarding the Tfh related immunological players that influence Dengue vaccine responses as depicted in **Fig 7**: Right from the first step of innate sensing of incoming Dengue virus by innate cells such as monocytes, we believe that micro-environmental changes occur leading to the differentiation of Tfh cells which subsequently increases in the peripheral detection of CXCL13/BLC. Tfh cells eventually helps in the recruitment of B cells and the formation of the GC that favors the production of high affinity antibodies and antibodies that have virus neutralization functionality. Circulating Tfh, CXCL13/BLC, and Innate cells could ultimately influence favorable vaccine responses, which potentially explains the dengue serostatus effect.

**Fig 7:**
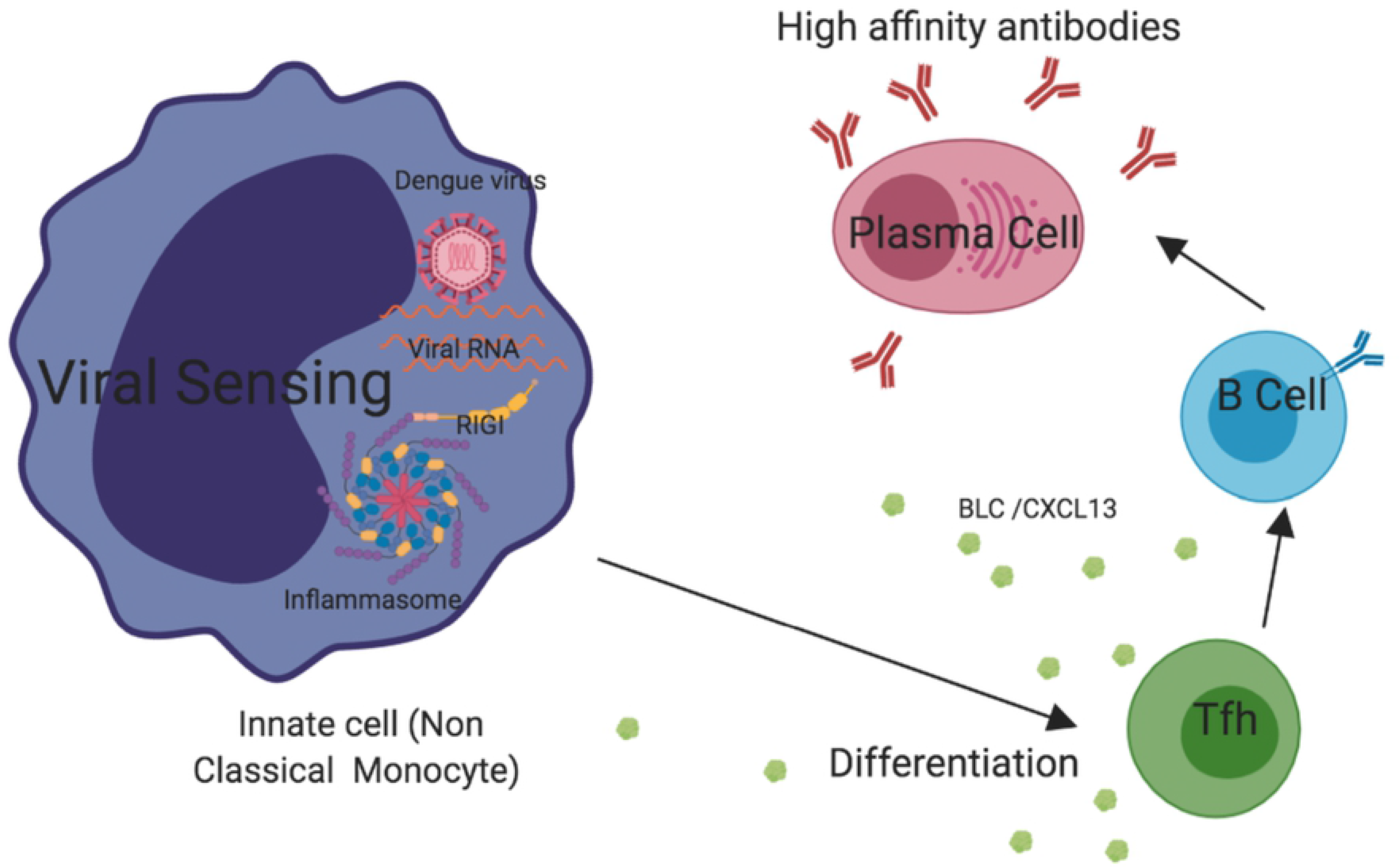
Proposed model. Innate cells such as monocytes are the primary targets of the dengue virus. They sense the virus leading to micro-environmental changes that help the differentiation of Tfh cells. Tfh cells in turn, secrete analytes such as BLC/CXCL13, which help in the recruitment of B cells and the formation of the GC that favors the production of high affinity antibodies and antibodies that have virus neutralization functionality.

## Material and methods

### Vaccination (TV003) and Subjects (Cohort)

TV003, is one of the tetravalent formulations selected by Butantan Institute for further development and manufacture as a lyophilized tetravalent Dengue vaccine (Butantan-DENV). TV003 is a live-attenuated vaccine, which contains all four Dengue serotypes (DENV1–4). Subjects enrolled in this trial were either immune or naïve to Dengue (seropositive or seronegative). More details about the vaccine and the cohort could be found in the referenced study of the Phase II trial (53). This study was approved by the local ethics committee of the Medical School of the University of São Paulo, the Brazilian Research National Ethics Council, the Brazilian National Technical Biosafety Commission, and the Brazilian Health Regulatory Agency. The study was done in accordance with the Declaration of Helsinki, International Conference of Harmonization guidelines, and Good Clinical Practice guidelines. All patients provided written informed consent. An independent data safety monitoring board oversaw the study. This trial is registered with ClinicalTrials.gov, NCT01696422.

### Neutralizing antibody titers

Neutralizing antibodies to DENV were determined by plaque-reduction neutralization titer (PRNT) assays as previously described (53). Briefly, for all doners, an initial serum dilution of 1/5 was used for PRNT_50_ assays. Seropositivity was defined by PRNT_50_ cutoff titers (≥1/10) before immunization and at any timepoint up to 91 days after vaccination (day 28, 56, or 91). For DENV-naive participants, seroconversion was defined by PRNT_50_ cutoff titers (≥1/10). For participants previously exposed to DENV, seroconversion was defined as a four-fold or higher increase in pre-existing neutralizing antibody titer after immunization (53).

### Flow cytometry and antibodies

PBMCs from Dengue seropositive and Dengue seronegative volunteers at day 0 were incubated with fluorochrome-conjugated antibodies for at least 15–20 min at 4 °C or on ice, protected from light. The following fluorochrome-conjugated anti-human antibodies were used: CD3 (HIT3α) (dilution: 1/100; Cat. Number: 300324), CD4 (RPA-T4) (dilution: 1/100; Cat. Number: 300518), CD25 (BC96) (dilution: 1/100; Cat. Number: 302608), CD38 (HIT2) (dilution: 1/33; Cat. Number: 303532), PD-1 (EH12.2H7) (dilution: 1/20; Cat. Number: 329918), CXCR3 (G025H7) (dilution: 1/20; Cat. Number: 353716), CCR6 (G034E3) (dilution: 1/20; Cat. Number: 353426), CXCR5 (J252D4) (dilution: 1/100; Cat. Number: 356908), ICOS (C398.4A) (dilution: 1/50; Cat. Number: 313518), KI-67 (KI-67) (dilution: 1/20; Cat. Number: 350504), CD16 (3G8) (dilution: 1/50; Cat. Number: 302025), CD1c (L161) (dilution: 1/50; Cat. Number: 331519), CD163 (GHI/G1) (dilution: 1/25; Cat. Number: 333608), CD11c (3.9) (dilution: 1/50; Cat. Number: 301636), HLA-DR (L243) (dilution: 1/50; Cat. Number: 307650), CD80 (2D10) (dilution: 1/20; Cat. Number: 305210), CD14 (M5E2) (dilution: 1/25; Cat. Number: 301852), CX3CR1 (2A9-1) (dilution: 1/25; Cat. Number: 341604), CD303 (201A) (dilution: 1/50; Cat. Number: 354212), PD-L1 (29E,2A3) (dilution: 1/20; Cat. Number: 329708), CD56 (HCD56) (dilution: 1/20; Cat. Number: 318318), CD3 (HIT3a) (dilution: 1/100; Cat. Number: 300316), CD19 (HIB19) (dilution: 1/100; Cat. Number: 302216), CD19 (HIB19) (dilution: 1/100; Cat. Number: 302252), CD38 (HIT2) (dilution: 1/50; Cat. Number: 303524), CD10 (HI10a) (dilution: 1/50; Cat. Number: 312212), IgD (IA6-2) (dilution: 1/50; Cat. Number: 348208), CD20 (2H7) (dilution: 1/25; Cat. Number: 302304), KI-67 (KI-67) (dilution: 1/20; Cat. Number: 350506), BCL-2 (100) (dilution: 1/100; Cat. Number: 658606), were all from BioLegend. IgM (G20-127) (dilution: 1/20; Cat. Number: 551079), CD21 (B-ly4) (dilution: 1/50; Cat. Number: 555422), IgG (G18-145) (dilution: 1/20; Cat. Number: 561298),CD21 (B-ly4) (dilution: 1/50; Cat. Number: 557327), were from BD Biosciences. CD141 (AD5-14H12) (dilution: 1/50; Cat. Number: 130-090-51) was from Miltenyi. FOXP3 (PCH1.1) (dilution: 1/20; Cat. Number: 53-4776-42) was from Thermo Fisher eBioscience. CD45RA (2H4LDH11LDB9) (dilution: 1/100; Cat. Number: IM2711U) was from Beckman Coulter. LIVE/DEAD Fixable Dead Cell Stain (Life Technologies) (dilution: 1/ 100; Cat. Number: L34957) was used to gate on live cells. Samples were acquired on a BD LSR II.

### Cytokine and chemokine analysis

Donor plasma was analyzed for chemokine/cytokine levels using ProcartaPlex (ThermoFisher Scientific). The following human premixed chemokine/cytokine panels was used: Cortisol, BLC, CD30L, CD40-L, C-TACK, D-Dimer, EDA-1, ENA78, Eotaxin, Eotaxin-2, FGF-23, Fibrinogen, GDF-15, Ghrelin, Granzyme B, HGF, HVEM, IL-1beta, IL-16, IL-18, IL-33, IL-37, IP-10, MCP1, MDC/CCL22, MIF, MMP-1, Nectin-2 (CD112), NT5E (CD73), OPG, PD-1, Perforin, sFas-Ligand, TIM-3, TNFRI, TNFRII, Trail-R1, TSLP, Tweak, ULBP-1, ULBP-3, and VEGF-R3. The manufacturer’s protocol was followed. Data were acquired on a Bio-Plex 200 System (using bead regions defined in thermo fisher Scientific protocol) and analyzed with the Bio-Plex Manager 6.1 software from Bio-Rad or a Luminex™ FLEXMAP 3D™ System using bead regions defined in the protocol and analyzed using Belysa Curve Fitting Software (Sigma Aldrich). Standard curves were generated, and sample concentrations were calculated in pg/mL.

### AUCp Calculation and Breadth

In summary, we measured the log-transformed Area Under the nAb titer Curve pre boost (AUCp) using the trapezoid rule in R. Breadth was calculated on a binary basis of detection / absence (nAb >10) for each serotype for at least one timepoint, and then summed up across serotypes, as per the clinical definition (Breadth 0-4) (53). The outcome was then dichotomized into high and low breadth groups, where high corresponds to a detectable nAb response to all 4 serotypes, and low to values under 4.

### t-distributed stochastic neighbor embedding (t-SNE) Flow cytometry

Individual FCS files generated by the BD FACS Diva software were imported into FlowJo Software for pre-gating on CD4+ live cells for the T cell panel and live cell cells for the innate panel. Selected events were then exported: from which an identical number of events per participant were then randomly subsampled into R. For bioinformatic analysis of flow cytometry data, a custom script was made for unsupervised clustering of cells based on similarity of their marker expression.

Batch correction was applied to account of variations across experiments, using the following methodology: a spline-model was built for each fluorescent marker on the quantile transformed data of a selected representative batch (oe experiment). These models were then used to correct fluorescence values of quantile transformed expression for other batches. Control “bridge” samples were leveraged to assess for tSNE overlap following batch correction as a measure of accuracy. Following batch correction, dimension reduction (PCA followed by tSNE) was performed, followed by RPhenograph clustering using the louvain method.

Cluster profiles are represented using the min-maxed transformed Median Fluorescence Intensity (MFI) for each marker on a linear transformation as a heatmap. Cell frequencies per cluster per participants were then compared across discrete outcomes (pre-vaccination serostatus and Breadth) using a Welch T-test, or were correlated with serotype-specific AUCp outcomes using Spearman correlation.

## Acknowledgments

We would like to acknowledge King Abdulaziz University and their fellowship program for their support. Also we would like to acknowledge Dr. Marita Chakhtoura and Dr. Roshell Muir for technical assistance and the constructive discussions. Finally, we would like to thank the entire study participant who made this study possible.

## Supporting information

**S1 Fig:**
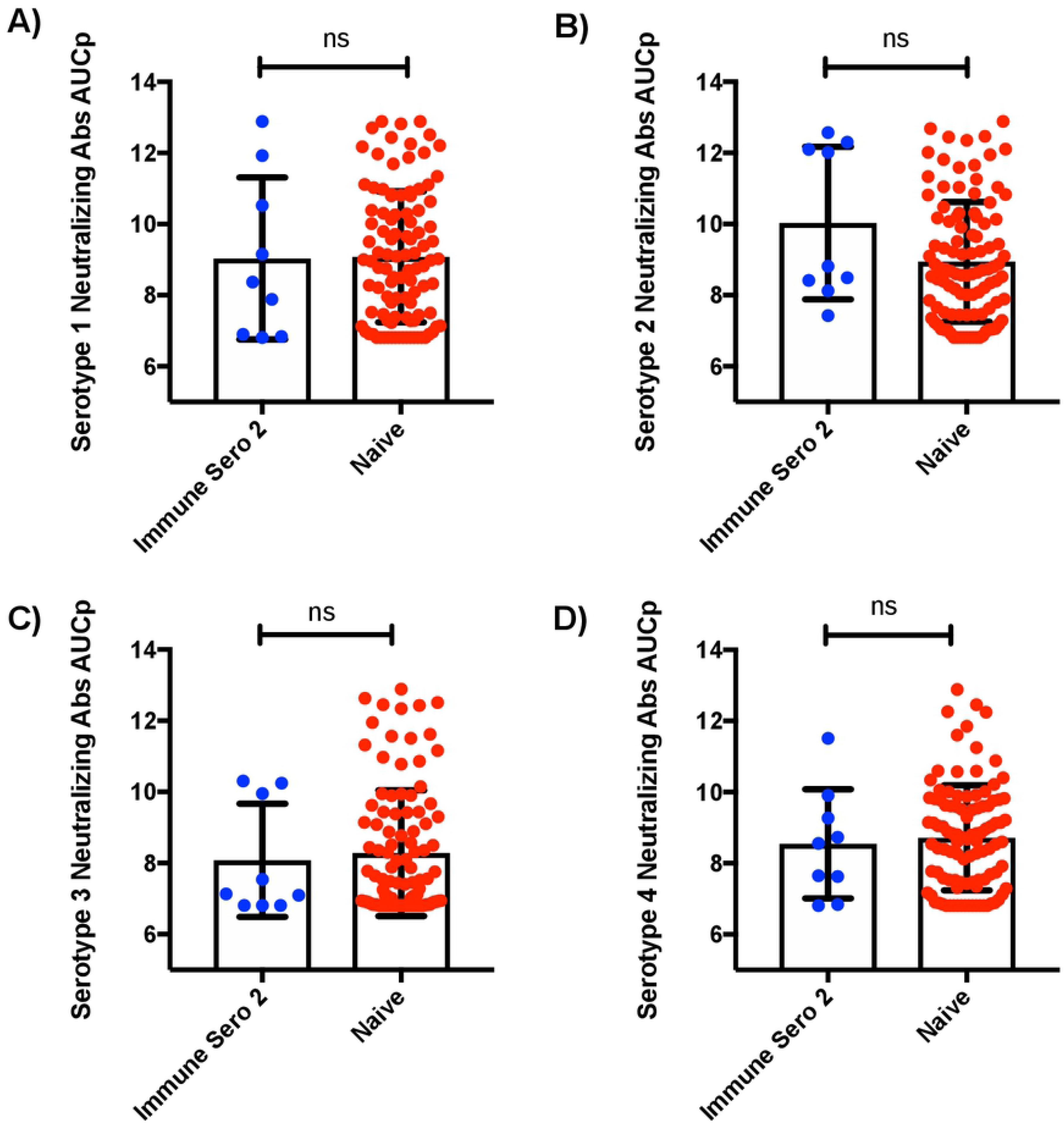
Pre-existing immunity to serotype 2 only does not increase dengue serotype 2 neutralizing antibody titers (AUCp). **(A)** Y axis is Dengue serotype 1 neutralizing antibody titers (AUCp). **(B)** Y axis is Dengue serotype 2 neutralizing antibody titers (AUCp). **(C)** Y axis is Dengue serotype 3 neutralizing antibody titers (AUCp). **(D)** Y axis is Dengue serotype 4 neutralizing antibody titers (AUCp). “Immune Sero 2” are subjects who only had Dengue Serotype 2 neutralizing antibody titers (AUCp) at pre-vaccination/baseline time point. Seronegative subjects with no existing antibodies to any of the four Dengue virus serotypes at pre-vaccination/baseline time point are referred to as “naïve”. This was accomplished using plaque reduction neutralization test (PRNT). Unpaired non-parametric Mann Whitney test 95% confidence was calculated for **(A-D)**.

**S2 Fig:**
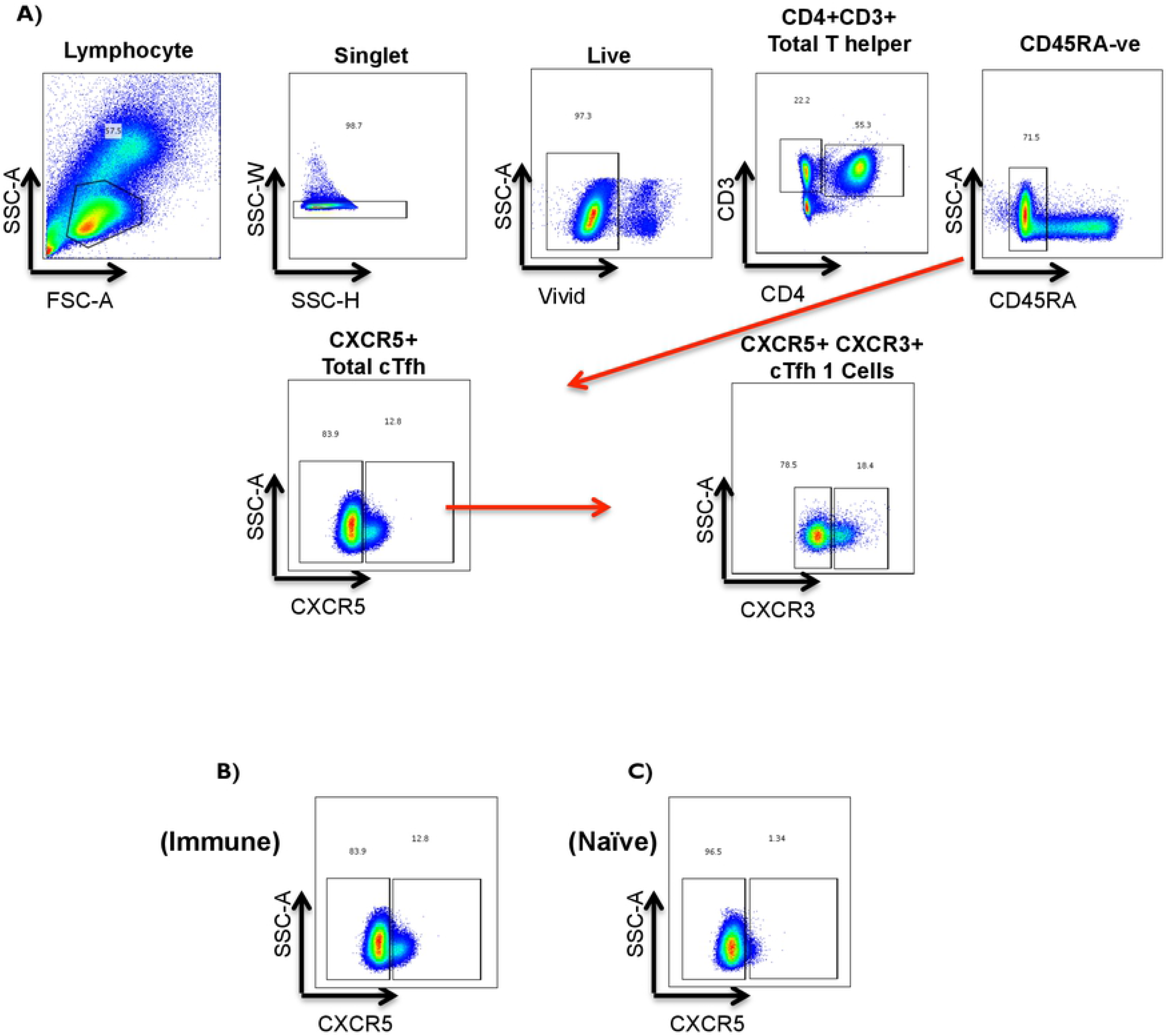
T Cell Panel Gating strategy. **(A)** Gating strategy of the T Cell Panel; **T Helper Cells:** CD3+ CD4+. Total **circulating T follicular Helper Cells (cTfh):** CD3+ CD4+ CD45RA- CXCR5+**. cTfh1:** CD3+ CD4+ CD45RA- CXCR5+ CXCR3+**. cTfh2,17:** CD3+ CD4+ CD45RA- CXCR5+ CXCR3- **. (B)** Immune subject example of total cTfh at pre-vaccination/baseline time point. **(C**) Naïve subject example of total cTfh at pre-vaccination/baseline time point. Before vaccination, seropositive subjects with existing antibodies to any of the four Dengue virus serotypes due to prior infection were referred to as (“Immune”) or seronegative subjects with no existing antibodies to any of the four Dengue virus serotypes were referred to as (“Naïve”).

**S3 Fig:**
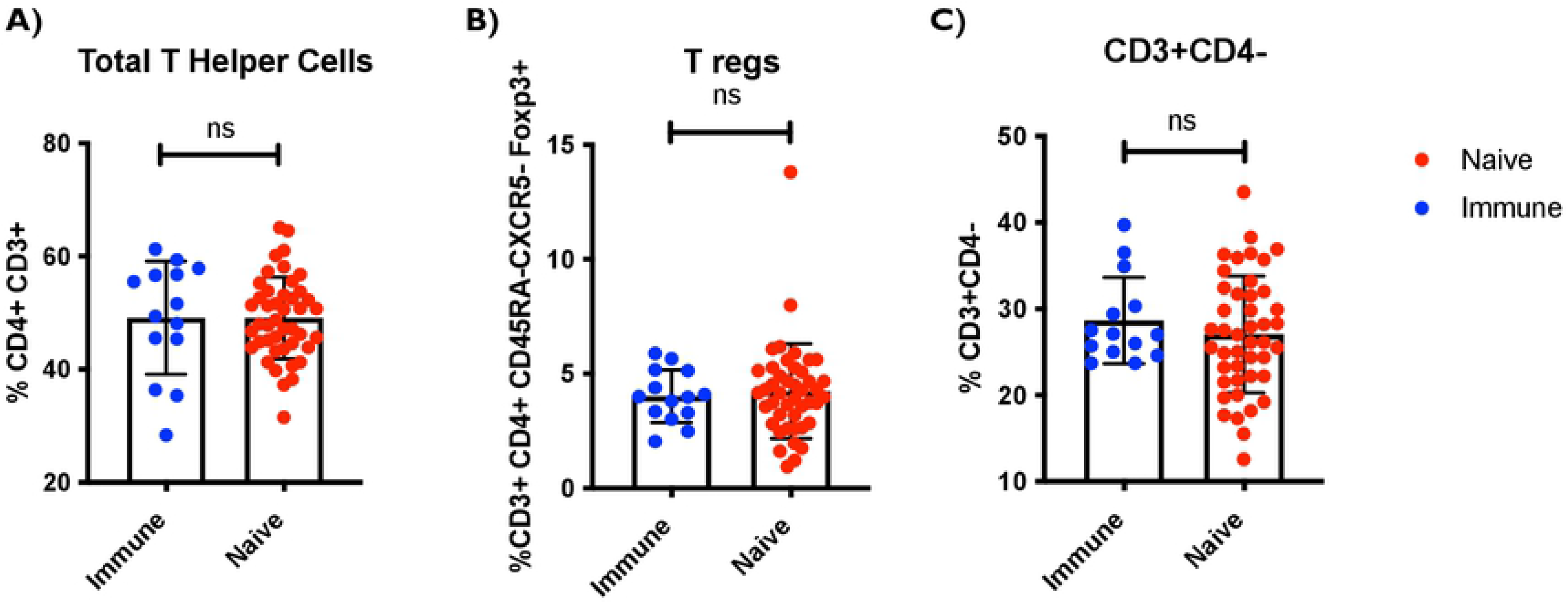
No difference at pre-vaccination frequency of major T cell subsets was observed between seropositive (Immune) and seronegative (Naïve) groups. **(A)** Y axis is the total T helper cell frequency CD3+ CD4+. **(B)** Y axis is the total Treg frequency CD3+ CD4+ CD45RA- CXCR5- Foxp3+. **(C)** Y axis is the total CD3+ CD4- frequency. Pre-vaccination/baseline subjects PBMCs and optimized T cell panel for ex-vivo staining was utilized. 1 million cells per panel were stained. Unpaired non-parametric Mann Whitney test 95% confidence was calculated for **(A-C). (**Naive N=44, Immune N=14). Before vaccination, Seropositive subjects with existing antibodies to any of the four Dengue virus serotypes due to prior infection were referred to as (“Immune”) or seronegative subjects with no existing antibodies to any of the four Dengue virus serotypes were referred to as (“Naïve”).

**S4 Fig:**
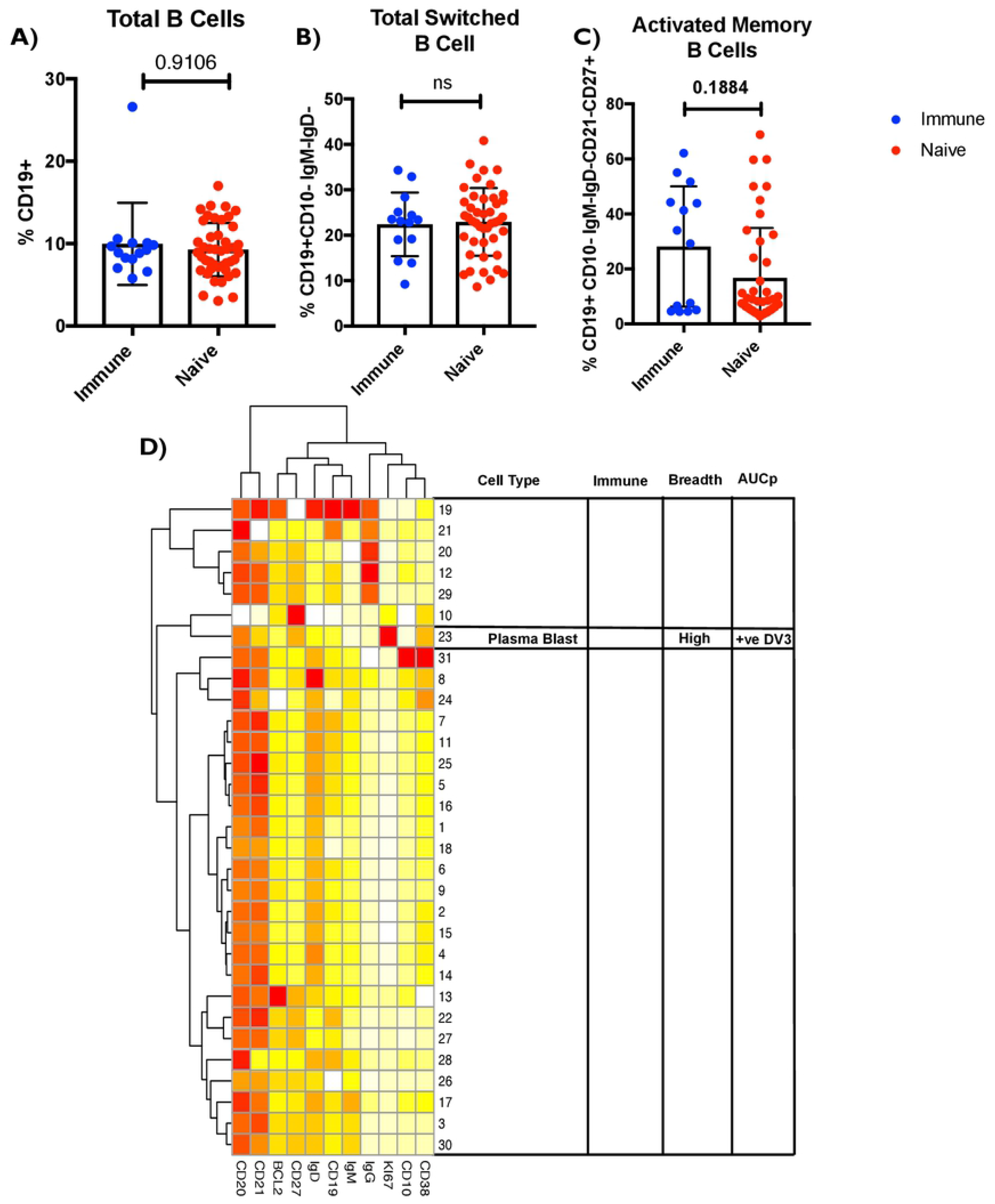
No difference at pre-vaccination frequency of major B cell subsets was observed between seropositive (Immune) and seronegative (Naïve) groups. **(A)** Y axis is the total B cell frequency: CD19+. **(B)** Y axis is the total switched B cell frequency: CD19+CD10-IgM-IgD-. **(C)** Y axis is the activated memory B cell frequency: CD19+CD10-IgM-IgD-CD21-CD27+. Pre-vaccination/baseline subjects PBMCs and optimized B cell panel for ex-vivo staining was utilized. 1 million cells per panel were stained. **D)** Heatmap of the 31 different clusters expressing different phenotypes using the optimized B cell panel with markers are shown in the x axis. Among them is cluster 23 that represents a B cell plasmablast cluster. The table identifies cell type, immune group pre-vaccination/baseline difference, breadth and neutralizing antibody titers (AUCp). Cells were pre gated on live cells and CD19, the expression was scaled and used to perform clustering which was visualized using t-Distributed Stochastic Neighbor Embedding (tSNE). Cell frequency per cluster/ per sample was derived and used for statistical testing between high and low breadth across donors. Subjects who mounted neutralizing antibodies against all four serotypes (termed “high” breadth) were compared to subjects who mounted neutralizing antibodies against < 3 serotypes (termed “low” breath). Pre-vaccination/baseline subject samples were associated with breadth of the response and neutralizing antibody titers (AUCp) post vaccination. (Naïve n=44 Immune n=14). Before vaccination, seropositive subjects with existing antibodies to any of the four Dengue virus serotypes due to prior were referred to as *“Immune”) or seronegative subjects with no existing antibodies to any of the four Dengue virus serotypes were referred to as (“Naïve”). Unpaired non-parametric Mann Whitney test 95% confidence was calculated for **(A-C)**. Non-parametric Spearman Correlation with 95% confidence intervals was used for correlations with neutralizing antibody titers (AUCp column).

## Notes

### Competing Interest Statement

The authors have declared no competing interest.

